# Effects of multiple cell regulators on curli gene expression in *Escherichia coli*

**DOI:** 10.1101/2025.07.09.663852

**Authors:** Maryia Ratnikava, Olga Lamprecht, Victor Sourjik

**Affiliations:** Max Planck Institute for Terrestrial Microbiology and Center for Synthetic Microbiology (SYNMIKRO), Karl-von-Frisch Strasse 14, 35043 Marburg, Germany

**Keywords:** curli, transcriptional control, c-di-GMP, motility, RpoS, bimodality

## Abstract

Curli amyloid fibers is the key protenacious component of extracellular matrix in *Escherichia coli*. Regulation of curli expression is highly complex and depends on the cellular response to diverse environmental conditions. Although many genetic determinants of curli production are known, previous studies were performed under diverse conditions and using different strains of *E. coli* or *Salmonella*. Furthermore, the origin of bimodal curli expression in bacterial population remains unknown. Here, we systematically investigated the role of multiple cell factors in expression of curli structural genes *csgBA* in unstructured planktonic *E. coli* culture. We observed that multiple regulators involved in regulation of stress response, cell motility, cell physiology and metabolism, as well as in formation of extracellular matrix and maintenance of DNA architecture modulated expression of curli by either promoting or repressing activity of *csgBA*. We further elucidated which regulators act upstream of the master transcription factor CsgD and which are crucial for bimodality of curli gene expression. We also investigated the impact of individual diguanylate cyclases on the regulation of the *csgBA* activity. This study provides an overview of regulation of curli gene expression in planktonic *E. coli* culture in the absence of any microenvironmental gradients.

## Introduction

Most bacteria are able to produce extracellular matrix that promotes formation of multicellular structures, giving protection against adverse environmental factors [1–3]. Curli are the major protenacious component of the extracellular matrix produced by *Escherichia coli* [4–8]. They are highly stable to degradation, thus supporting environmental cell persistence and cell stress survival [5, 6, 9]. The formation of curli is known to be triggered by a combination of suboptimal growth conditions and environmental stresses, such as slow growth, nutrient starvation, low temperatures, low osmolarity etc [2, 10–15].

In *E. coli*, genes required for curli formation are organized in two oppositely transcribed operons *csgBAC* and *csgDEFG* [2, 12, 15]. They are separated by a 754 bp long intergenic non-coding region, which is one of the largest and most heavily regulated in *E. coli* [5, 16], with curli expression dependent on the cellular response to multiple environmental conditions [8, 11, 13, 15, 17]. The *csgBAC* operon controls production of curli fibers, while the *csgDEFG* operon encodes a master curli regulator CsgD and accessory proteins required for assembly and secretion of the curli structural subunits CsgB and CsgA [5, 9, 18, 19]. CsgD activates expression of curli structural genes as well as its own expression by binding to multiple sites within the *csg* region [9, 20]. The expression of curli genes depends on the general stress response regulator RpoS and it is further known to be affected by multiple transcriptional factors, small RNAs (sRNAs) and second messenger bis-(3′–5′)-cyclic di-GMP (c-di-GMP) [2, 5, 14, 15, 17, 21–31]. A number of transcription factors have been reported to bind within the *csg* region and collaboratively modulate expression of curli genes from σ^70^-and σ^S^-dependent promoters [16, 23, 27, 32, 33], including those associated with metabolic transitions, two-component signaling and flagellar gene expression, as well as regulators maintaining DNA architecture (IHF and H-NS) [2, 8, 16, 33–37].

Intracellular c-di-GMP concentration, another key regulator of curli gene expression, is itself under complex control by a set of diguanylate cyclases (DGCs) and phosphodiesterases (PDEs) [29, 38, 39]. In total, 14 DGCs have been identified in non-pathogenic *E. coli* strains, including two degenerate proteins that lack the catalytic function [39, 40]. Curli gene expression is primarily modulated by DgcE and DgcM [28, 29, 31, 41, 42]. Elevated c-di-GMP levels promote expression of curli by derepressing MlrA, the transcriptional activator of *csgD* [28, 31]. Other active DGCs, including DgcC, DgcP, and the degenerate DGC CsrD, have been shown to further regulate CsgD activity in *E. coli* [39].

Expression of curli within a population is restricted to a group of cells, resulting in formation of curli-producing and non-producing cells [43–47]. Heterogeneous production of curli in macrocolony and submerged biofilms could be driven by presence of microenvironmental and metabolic gradients [43, 48–50]. However, bimodal curli expression is also observed in an isogenic well-stirred planktonic culture of *E. coli* [43, 46], suggesting that bimodality is an inherent property of curli gene activation.

Here, we systematically assessed how multiple cellular regulators and DGCs contribute to expression of the *csgBA* genes in unstructured planktonic *E. coli* culture grown under continuous mixing and thus in the absence of any gradients or biofilm formation. We confirmed involvement of several known curli regulators and demonstrated impact of several regulators that were not previously shown to control curli expression. In contrast, other previously proposed regulators of curli expression showed no impact under our growth conditions. Furthermore, we classified which regulators act upstream of *csgD* and uncovered a set of regulators crucial for bimodality of expression of curli structural genes in absence of microenvironmental gradients. Our study provides a systematic overview of curli regulation in unstructured planktonic *E. coli* culture.

## Materials and methods

### Bacterial strains and plasmids

All strains and plasmids used in this study are listed in S1 Table. An RpoS^+^ derivative of *E. coli* W3110 [51] that was engineered to encode a chromosomal transcriptional superfolder green fluorescent protein (sfGFP) reporter downstream of the *csgA* gene (VS1146) [44] was used as the wild-type (WT) strain.

Majority of gene deletions were obtained with the help of P1 phage transduction as described in [52] with some modifications. Strains of the Keio collection [53] were used as donors for the transfer of the kanamycin resistance cassette, which disrupted the coding sequence leaving intact only start codon and last 21 nucleotides (except for *dgcO* deletion, where first 342 and last 21 nucleotides were retained). Deletion of *dgcN* led to the in-frame addition of the 240 nucleotides encoding for a part of Tn5 transposase to the upstream located *yfiR* gene. In the case of *csgD*, the deletion strain was constructed by lambda red recombination method, with first 200 nucleotides and last 30 nucleotides being kept to avoid possible disruption of the common curli gene regulatory region. In the strains that showed differences in curli expression from WT (except for Δ*ihfA*), the kanamycin resistance cassette was eliminated using FLP recombinase as described in [54] to confirm absence of polar effects of the cassette [55]. Obtained gene deletions were confirmed by PCR with primers flanking the corresponding genes followed by Senger sequencing.

For complementation analysis of curli-expression deficient strains, *csgD* gene was cloned into the pTrc99A vector by restriction-ligation [56] and transformed in cells using TSS method as described in [57] with modifications.

### Growth conditions of cultures

To analyze *csgBA* expression levels and fraction of curli positive cells by flow cytometry approach, *E. coli* cultures were inoculated from lysogeny broth (LB) agar plates (10 g tryptone, 5 g yeast extract, 5 g NaCl, 15g Bacto Agar per liter; pH 7.0) supplemented with 50 µg/ml kanamycin where necessary and grown in 5 ml of LB medium at 37°C at 200 rpm in 100 ml flasks in a rotary shaker overnight (14-16 h). Obtained cultures were diluted 1:100 in 5 ml of tryptone broth (TB) medium (10 g tryptone, 5 g NaCl per liter; pH 7.0) and grown at 30°C in a rotary shaker at 200 rpm in 100 ml flasks for a time required to detect maximum *csgBA* expression levels (appr. 12-34 h depending on the strain).

To analyze *csgBA* expression levels and growth curves (OD_600_) by plate reader analysis, *E. coli* cultures were inoculated from LB agar plates supplemented with 50 µg/ml kanamycin where necessary and grown in liquid LB at 37°C at 200 rpm in 100 ml flasks in a rotary shaker overnight (14-16 h). Obtained cultures were diluted 1:100 in TB medium and incubated at 30°C in INFINITE 200 PROplate reader (Tecan Group Ltd., Switzerland) for 24-25 h.

For complementation analysis, overnight cultures were inoculated from LB agar plates supplemented with 100 µg/ml ampicillin and grown in 5 ml of TB supplemented with 100 µg/ml ampicillin and indicated levels of inducer isopropyl β-d-1-thiogalactopyranoside (IPTG) at 30°C at 200 rpm in 100 ml flasks in a rotary shaker overnight (18-26 h).

### Flow cytometry analysis

Flow cytometry method was used for sfGFP expression measurements. Aliquots of 40-70 µl of bacterial cultures were mixed with 2 ml of PBS (0.14 mM NaCl, 2.7 mM KCl, 1.5 mM KH_2_PO_4_, 8.1 mM Na_2_HPO_4_ per liter; pH 7.0) and obtained samples were subjected to the flow cytometry analysis within next 30 min. Before measurements the probes were vigorously vortexed to disrupt cell clumps and then immediately subjected to BD LSRFortessa Sorp cell analyzer (BD Biosciences, Germany) using 488-nm laser. 50 000 individual cells were analyzed in each experimental run. Recorded events were further filtered to exclude cell duplets and debris with the help of forward scatter (FSC) and side scatter (SSC) plots (SSC-H vs. SSC-W and FSC-A vs. SSC-A). Data were analyzed using FlowJo software version v10.7.1 (FlowJo LLC, Ashland, OR, US). The gate used to differentiate subpopulation of negative cells from curli-positive was defined as corresponding fraction of curli-negative cells in W3110 RpoS+ strain carrying no *csgBA* genomic reporter.

### Statistical analysis

Expression of *csgBA* genes and fraction of curli-positive cells were statistically analyzed using ordinary one-way ANOVA followed by Dunnett’s test to run multiple comparisons between WT and deletion strains. The significance level (α) was set to 0.05. The adjusted p-value between 0.01 and 0.05 is indicated with one asterisk (*), between 0.01 and 0.001 with two asterisks (**), between 0.001 and 0.0001 with three asterisks (***), p < 0.0001 with four asterisks (****). The statistical analysis was performed in GraphPad Prism version 10.0.0 for Windows (GraphPad Software, Boston, Massachusetts USA).

## Results

### Deletions of multiple regulator genes affect expression of curli fibers

In order to systematically investigate the role of different cellular factors in the regulation of curli structural genes in *E. coli*, we utilized a recently described strain that encodes a chromosomal GFP reporter as a part of the *csgBA* operon [44, 46]. We first constructed a library of 32 mutants in this strain, each carrying a deletion of a single gene selected based on its known involvement into extracellular matrix formation, control of cell stress response, metabolic transitions as well as maintenance of DNA architecture and cross-regulation between motility and curli production. We next measured activity of the *csgBA* reporter in planktonic cultures of these mutants using flow cytometry. Consistent with previous findings, expression of the *csgBA* genes was bimodal, with only a fraction of cells (~80%) activating GFP reporter in the overnight culture of the wild-type (WT) strain background (Fig. 1A) [46]. Half of the obtained deletion mutants showed the *csgBA* expression similar to that of WT (Fig. 1B-C and Fig. S1). Notably, these also included genes encoding regulators that were previously suggested by *in vivo* and *in vitro* experiments to bind to the *csg* intergenic region and/or modulate curli expression, either positively (*basR, bolA, rcdA, rstA*) [16, 58–60] or negatively (*btsR, mqsR, cpxR, rstA*) [16, 33, 35, 36, 61–63]. Similarly, deletion of *ariR, fnr, phoP* and *gadE* had no impact on the average *csgBA* expression levels (Fig. 1B), although they led to a moderate decrease of number of curli-expressing cells, whereas *gadE* – to a slight increase (Fig. S2).

**Figure 1.**
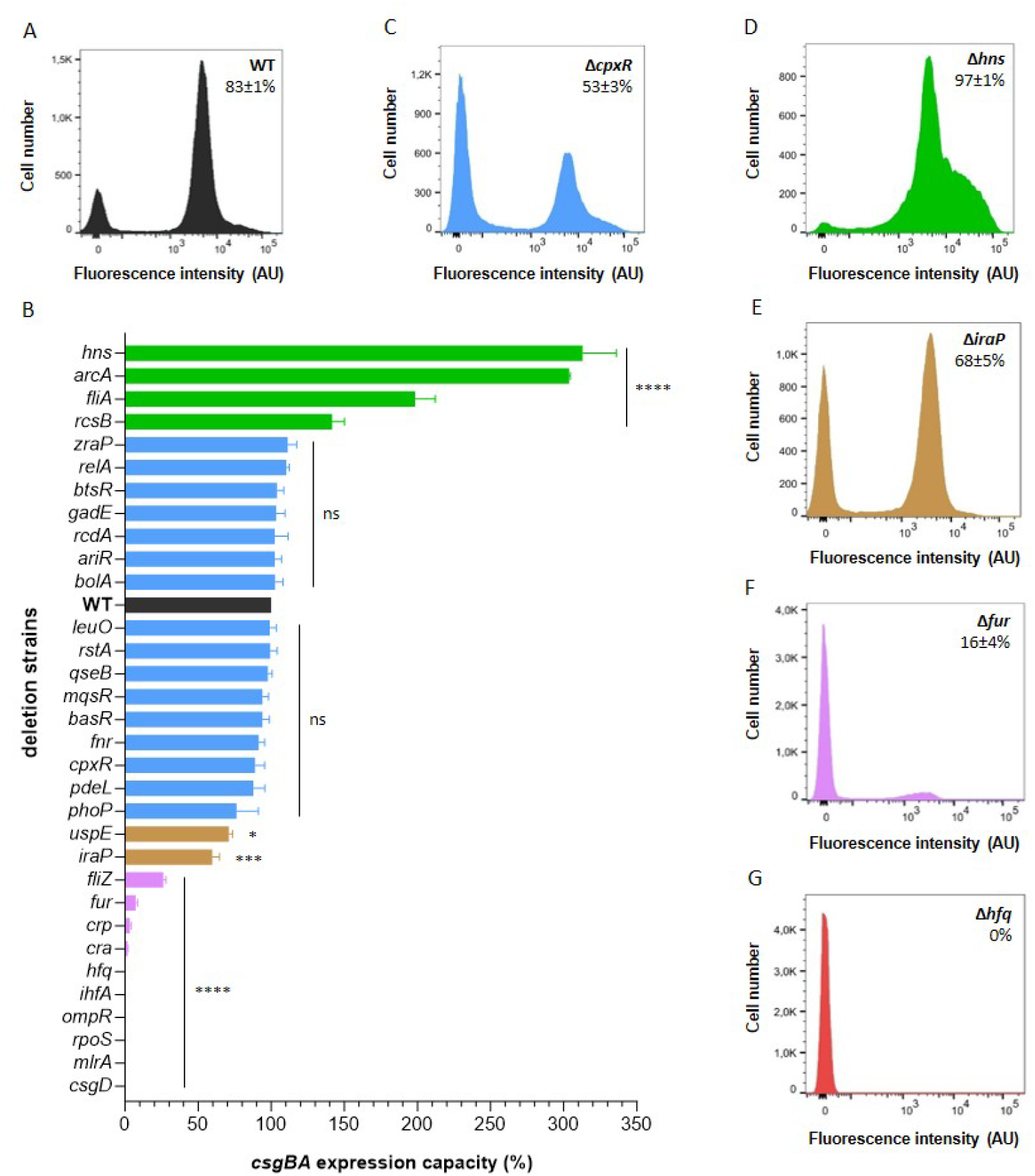
Curli gene expression responds differently to removal of individual genes. *E. coli* cells carrying genomic transcriptional reporter of the *csgBA* operon were grown in flasks with tryptone broth (TB) at 30°C under constant shaking until they reach maximal *csgBA* reporter levels and then subjected to the flow cytometry analysis. **(A)** Distribution of single-cell fluorescence levels in the wild-type strain (WT) strain. **(B)** Activity of the *csgBA* transcriptional reporter in library of single deletion mutants compared to WT. WT is shown in black color and deletion strains with enhanced curli expression are indicated in green, with not affected – in blue, reduced – in brown, strongly impaired – in purple and entirely abolished – in red. Error bars indicate standard error of the mean (SEM) of at least 3 biological replicates. * at p = 0.01-0.05, ** at p = 0.01-0.001, *** at p = 0.001-0.0001, **** at p < 0.0001. **(C-G)** Distribution of single-cell fluorescence in indicated deletion strains with not affected **(C)**, enhanced **(D)**, reduced **(E)**, strongly impaired **(F)** and entirely abolished **(G)** *csgBA* expression. Fraction of positive cells in the population (mean of at least 3 biological replicates ± SD) is indicated for each strain. Note that the scale in the *y* axes is different for individual strains to improve readability.

Four mutants (Δ*hns*, Δ*arcA*, Δ*fliA* and Δ*rcsB*) showed increased activity of the *csgBA* transcriptional reporter as well as the number of curli-positive cells, with the effects of *hns* and *arcA* deletions being most pronounced (Fig. 1B, D and Fig. S2 and S3A).

Moderate reduction of the *csgBA* expression and the number of curli-expressing cells was observed in mutants lacking genes that encode stress response genes UspE and IraP, indicating positive but auxiliary involvement of these proteins in regulation of curli structural genes in a well-shaken *E. coli* planktonic culture (Fig. 1B, E and Fig. S2 and S3B). In contrast to the positive impact of *fliA* deletion, deletion of *fliZ* showed decreased activity of the *csgBA* reporter (Fig. 1B, S2, S3C). Deletion of genes encoding global regulators of carbon metabolism *cra* and *crp* and the regulator of iron homeostasis *fur* severely reduced expression of the *csgBA* operon (Fig. 1B, F and Fig. S2 and S3C).

Finally, expression of the *csgBA* genes was completely abolished by the removal of regulators known to directly bind to the *csg* intergenic region and be required for the activation of curli gene expression (*rpoS, mlrA, csgD, ompR* and *ihfA*)(Fig. 1B and Fig. S2 and S3D) [8]. Similar abolishment was observed upon disruption of *hfq* gene encoding a chaperone for sRNAs (Fig. 1B, G and Fig. S2).

### Effects of deletions of individual regulator genes on cell growth and dynamics of *csgBA* activation

In order to further explore how removal of individual regulators influences the dynamics of curli gene expression throughout the growth phase, we monitored cell growth and activity of the *csgBA* operon over time using plate reader. The observed *csgBA* expression levels were generally consistent with those measured in the batch culture experiment using flow cytometry (Fig. 2 and Fig. S4, S5, S6). An exception was *arcA* mutant, which showed either same as WT or less activation of curli fibers in the plate reader (Fig. 2B and Fig. S6B), likely because of its slower growth (Fig. 2A and Fig. S6A). Nonetheless, the deletion of *arcA*, and even more so of *hns* led to activation of *csgBA* expression at lower cell density, consistent with derepression of curli genes in these deletion strains. In contrast, removal of IraP – positive regulator of RpoS – delayed curli activation (Fig. 2 and Fig. S6). These observations were further confirmed by measuring reporter expression in corresponding strains grown in well-shaken planktonic cultures using flow cytometry (Fig. S7). Also in these measurements, curli gene expression in Δ*hns* strain was heterogeneous but not as clearly bimodal as in other strains.

**Figure 2.**
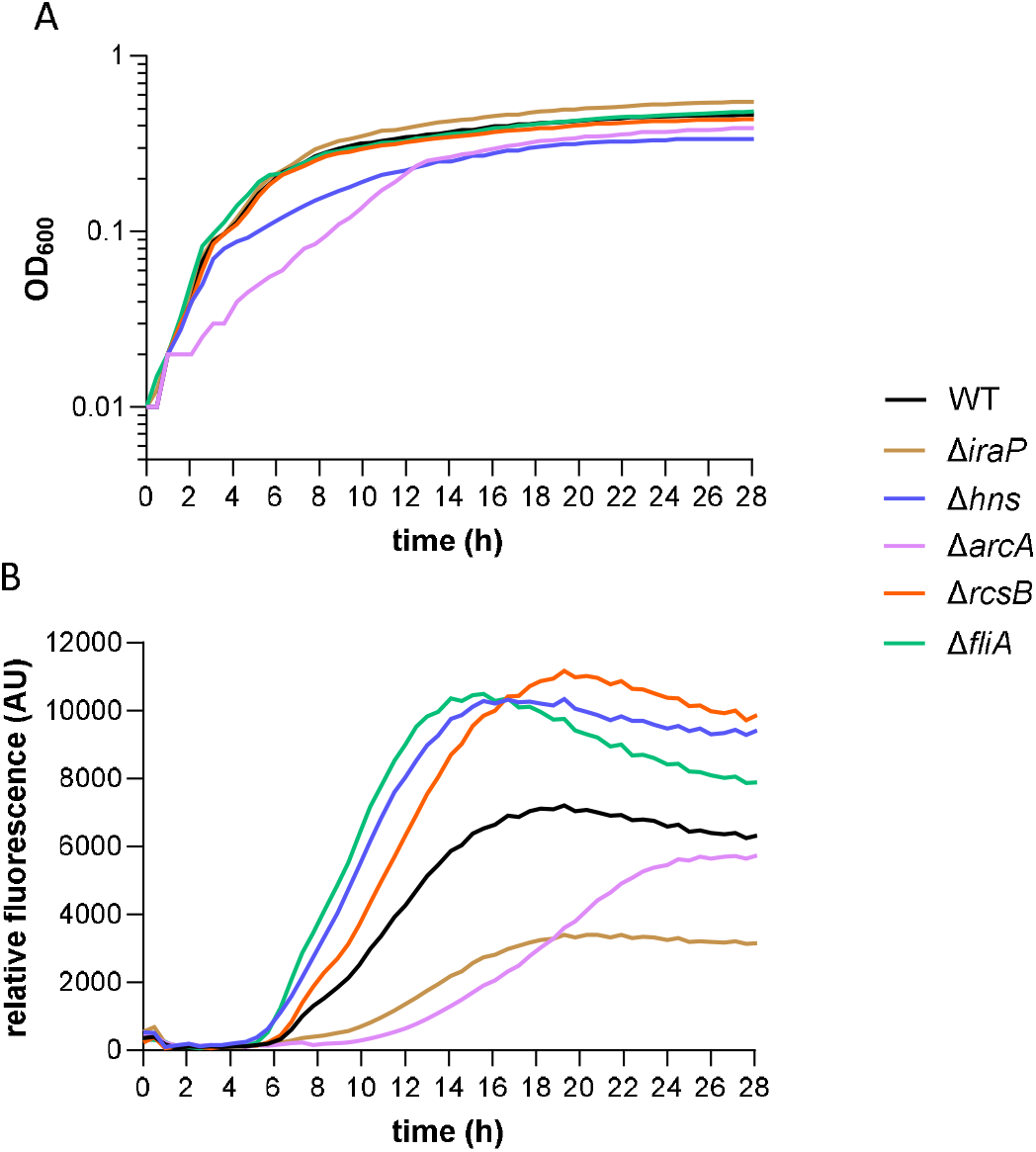
Removal of individual genes alters curli activation pattern. *E. coli* planktonic cultures carrying genomic transcriptional reporter of the *csgBA* operon were grown in a plate reader in TB at 30°C with shaking. Optical density (OD_600_) **(A)** and relative fluorescence (absolute fluorescence/OD_600_) **(B)** of WT and individual deletions strains during their growth in a plate reader. Shown graphs are representative of at least 3 independent replicates (other replicates are shown in Figure S6)

### Complementation of curli-expression deficient strains by ectopic expression of *csgD*

We further investigated whether the negative impact of dene deletions on the *csgBA* expression could be rescued by the expression of their immediate upstream regulator CsgD from a plasmid under control of the IPTG-inducible promoter. As expected, the expression of *csgD* at higher induction levels resulted in almost all cells expressing curli in the WT strain (Fig.3A). Restoration of the *csgBA* expression was observed in *fur, crp, cra* and *hfq* deletion strains, with formation of a well-distinguishable subpopulation of curli-positive cells at intermediate induction levels (Fig. 3B). Expression of CsgD in Δ*hfq* background also resulted in the increased number of curli-expressing cells, but with lower expression levels, and hence lesser separation between curli-positive and - negative cells. Our results thus suggest that these four regulators are primarily required for regulation upstream of *csgD* and are not essential for bimodal expression of curli fibers. CsgD could also activate *csgBA* expression in *rpoS* deletion strain (Fig. 3C), which confirms that the expression of curli structural genes can be induced from a vegetative promoter [12, 20, 32, 64–66]. Reduced separation between curli-positive and -negative cell subpopulations in this background at intermediate levels of CsgD induction indicates that the RpoS-dependent expression contributes to the bimodality of curli gene expression.

**Figure 3.**
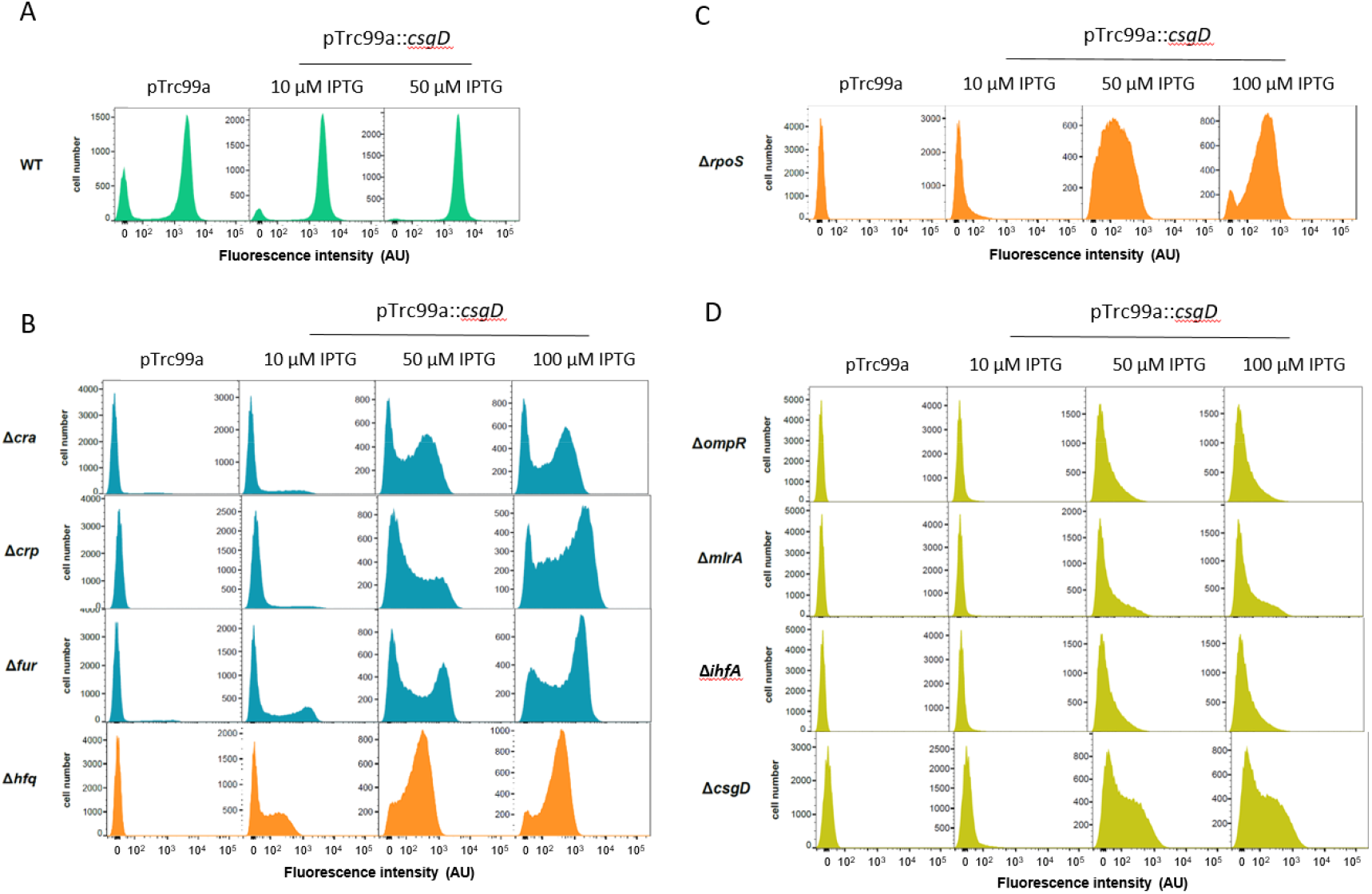
Expression of CsgD from a plasmid restores activity of *csgBA* operon to a different extent. *E. coli* cells carrying genomic transcriptional reporter of the *csgBA* operon and one of the indicated gene deletions were transformed with either empty pTrc99a plasmid (control) or with pTrc99a plasmid carrying *csgD* gene (pTrc99a::*csgD*). Expression from the vector was induced with 10, 50 and 100 μM IPTG. Bacteria were grown as well-shaked planktonic cultures in TB until stationary phase (OD_600_ ≥ 1.1) and cultures were subjected to the flow cytometry analysis. Distribution of single-cell fluorescence levels is shown. Note that the scale in the *y* axes is different for individual strains to improve readability.

In contrast, strains carrying deletions of *ihfA, ompR* or *mlrA* genes demonstrated only partial recovery of curli expression, without formation of a distinct positive population even at high IPTG induction levels (Fig. 3D). This suggests that these genes directly contribute to the regulation of curli gene expression downstream of CsgD and might modulate bimodal activation of the *csgBA* expression. Notably, although complementation by CsgD from the plasmid was better in the *csgD* deletion strain (lacking 421 nucleotides of the coding sequence (see Materials and Methods for details)), it failed to fully restore the *csgBA* expression, indicating that the native regulation of *csgD* and/or gene regulatory elements within its coding sequence might be required to fully induce expression of curli structural genes.

### Effects of deletions of individual DGC genes on curli expression

We next investigated how disruption of individual DGC-encoding genes affects *csgBA* expression in unstructured planktonic *E. coli* culture. Consistent with our previous study [46], deletions of the known curli regulators *dgcE* and *dgcM* nearly abolished curli expression (Fig. 4A and Fig. S8). Although removal of the majority of DGCs had no pronounced effects on the *csgBA* expression (Fig. 4A and Fig. S9A, S10A, S11), deletions of *dgcO, csrD* and *dgcC* genes resulted in markedly lower activity of the *csgBA* reporter and reduced number of cells expressing curli (Fig. 4A-D and Fig. S9A, S12), suggesting that only these DGCs contribute to curli regulation in *E. coli* in addition to DgcE and DgcM under conditions used in this study. Deletion of *dgcI* gene had a more complex effect, leading to decrease in the number of curli-expressing cells (Fig. S9A), as could be expected from removal of a DGC, but also resulting in a fraction of positive cells with highly elevated expression (Fig. 4B), and to transiently higher overall expression level of curli reporter in cell population in the early growth phase (Fig. 4C, D and Fig. S12).

**Figure 4.**
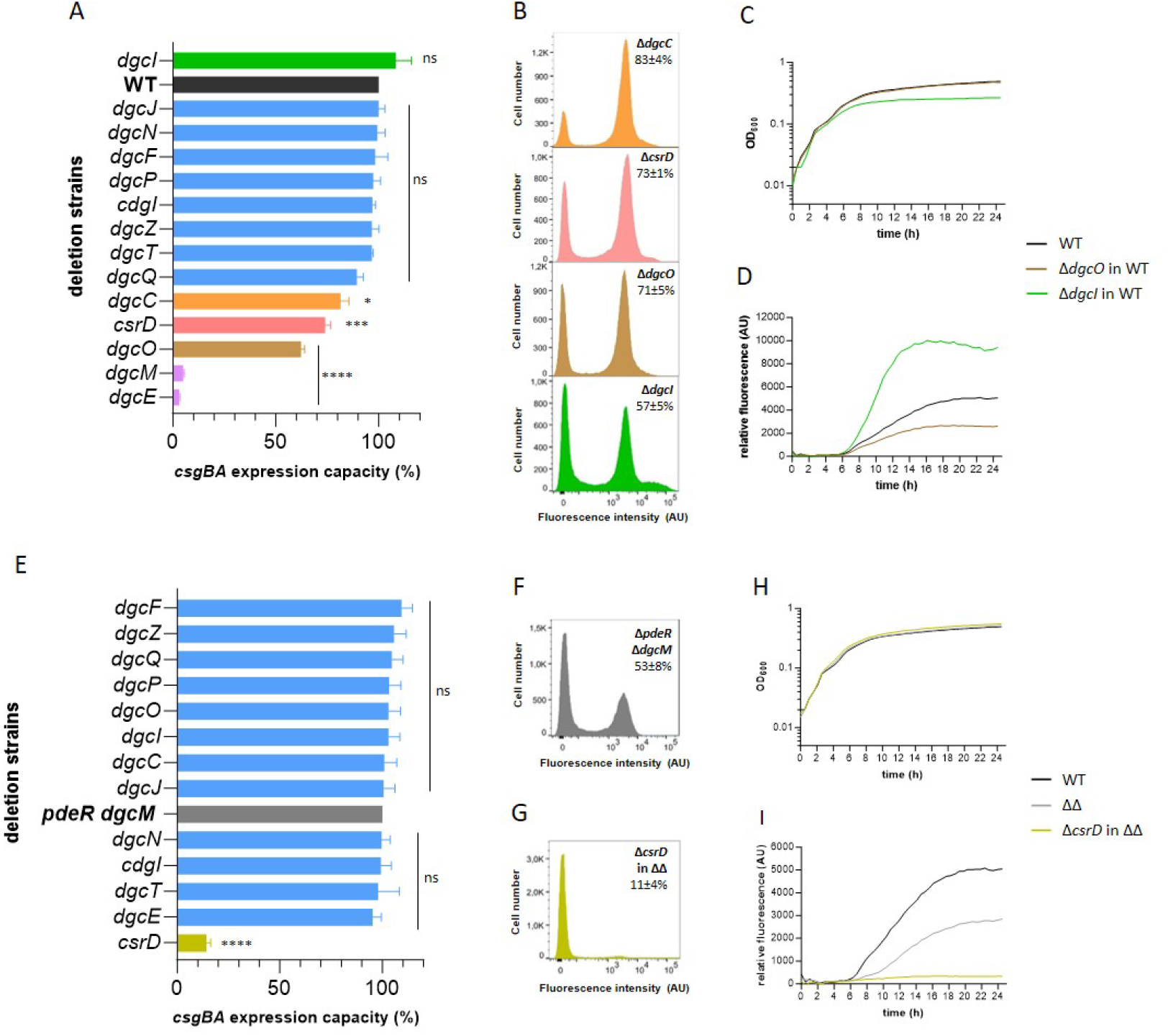
c-di-GMP regulatory network regulates curli expression in addition to DgcM/PdeR module. Activity of the *csgBA* transcriptional reporter **(A, E)** and distribution of single-cell fluorescence levels **(B, F, G)** in deletion strains lacking individual DGCs in WT background **(A, B)** and in absence of DgcM/PdeR regulatory module **(E-G)**. WT is shown in black color, strain lacking DgcM/PdeR regulatory module – in grey and deletion strains with not affected curli expression – in blue, with affected – in different colors. Error bars indicate SEM of at least 3 biological replicates. * at p = 0.01-0.05, ** at p = 0.01-0.001, *** at p = 0.001-0.0001, **** at p < 0.0001. *E. coli* cells carrying genomic transcriptional reporter of the *csgBA* operon were grown in flasks with TB at 30°C under constant shaking until they reach maximal *csgBA* reporter levels and then subjected to the flow cytometry analysis. Fraction of positive cells in the population (mean of at least 3 biological replicates ± SD) is indicated for each strain. Note that the scale in the *y* axes is different for individual strains to improve readability. **(C, H)** Optical density (OD_600_) and **(D, I)** relative fluorescence (absolute fluorescence/OD_600_) of WT and mutant strains lacking indicated DGCs during the growth in a plate reader (other replicates are shown in Figure S12, S13 and S14). *E. coli* cells carrying genomic transcriptional reporter of the *csgBA* operon were grown in TB at 30°C under constant shaking.

Since c-di-GMP is known to influence expression of the *csgBA* operon through the DgcM/PdeR-dependent *csgD* activation, we further measured reporter activity in triple mutants lacking DgcM/PdeR module and one of the individual DGCs (Fig. 4E). In agreement to our previous findings, activity of the *csgBA* operon was lower in a mutant lacking DgcM/PdeR regulatory module compared to WT but similarly bimodal (Fig. 4A, E, F, H, I and Fig. S13) [46]. Consistent with the expected dependence of c-di-GMP regulation on the DgcM/PdeR module, deletions of other cyclase genes had no effect in this background (Fig. 4E and Fig. S9B, S10B, S14), including the deletions of *dgcO, dgcI* and *dgcC*. In contrast, deletion of the *csrD* gene encoding a degenerate DGC led to a much stronger decrease of the reporter activity in the *dgcM/pdeR* knock-out background compared to the WT background (Fig. 4A, B, E-I and Fig. S9B, S15). This suggests that the expression of curli structural genes exhibits stronger dependence on CsrD in the absence of the local c-di-GMP regulatory module.

## Discussion

*E. coli* and other enterobacteria are known to express curli fibers in response to various environmental stresses. Multiple cellular factors were described to affect curli gene expression in structured biofilms communities, but the interpretation of previous studies was often complicated by environmental gradients in such communities. In this study, we systematically investigated the role of multiple cellular factors in regulation of curli gene expression under simpler conditions, in *E. coli* planktonic culture grown under continuous mixing.

Out of 32 tested strains, 21 individual gene deletions were found to modulate the *csgBA* expression levels and/or the number of curli-expressing cells in our experiments. Curli expression was elevated in several deletions strains, including those lacking components of the Rcs and ArcA/B signaling systems. The repression of curli expression by these pathways is in a good agreement with previous studies, with RcsB known to negatively regulate *csgD* expression [26, 30, 67–70], and ArcA known to repress transcription of *rpoS* [22, 70, 71]. An even more dramatic enhancement of curli gene expression was observed upon deletion of *hns*, confirming importance of this DNA-organizing factor as a curli repressor in *E. coli* [32]. Interestingly, H-NS was shown to activate curli expression in *S. typhimurium* but to repress in *E. coli*, by directly binding to the *csg* region and also by negatively affecting stability of *rpoS* mRNA [16, 24, 65, 72, 73].

In contrast, two stress regulators, the universal stress factor UspE and the RpoS-stabilizing factor IraP [74–77] showed positive impact on curli gene expression. The effect of IraP is likely mediated by the modulation of RpoS stability under starvation conditions [74]. Consistent with that, the deletion of *iraP* led to the delayed activation of the *csgBA* genes. We further observed the involvement of the iron-binding global transcriptional regulator Fur in positive curli regulation, which may be related to the previously observed transcriptional upregulation of *csgD* under iron limitation [78].

We further observed opposite effects of *fliZ* and *fliA* gene deletions, with curli gene expression being elevated in Δ*fliA* strain but lowered in Δ*fliZ* strain. The former effect was expected and it could be explained by the FliA-dependent activation of expression of *pdeH*, the major PDE of *E. coli* [79] that inhibits curli gene expression by lowering global levels of c-di-GMP [80].

However, FliZ was previously show to antagonize the activity of RpoS by binding within RpoS-dependent promoters [37, 79]. Why under our conditions FliZ has a positive rather than negative impact on curli expression remains to be investigated, but it might be due to its interplay with other regulators.

Dramatic reduction of curli expression was observed in the absence of global transcriptional factors Cra and CRP that are activated at low-nutrient conditions [11], thus confirming their importance for initiation of curli expression [8, 81–84]. Similar abortion of curli expression was observed upon deletion of *hfq*. Hfq is a chaperone that promotes pairing between sRNAs and their target mRNA, thus helping to regulate mRNA stability in response to different types of cell stress [85, 86]. The effect of *hfq* on curli gene expression is likely mediated by altered activity of sRNAs that are known to modulate curli expression, either directly by binding to the *csg* region [25, 26, 30, 69, 87, 88] or indirectly, through regulation of other cell factors that affect curli production, including RpoS [89–91]. Consistent with our observation, it has been previously shown that deletion of *hfq* leads to decreased *csgD* transcript levels in *S. enterica* [92]. Importantly, the defect of curli expression in all of *crp, cra, hfq* as well as *fur* deletion strains could be complemented by expressing CsgD from an inducible plasmid, confirming that these regulators act upstream of CsgD. However, the complementation was only partial in the case of *hfq* strain, indicating that a CsgD-independent impact of sRNA regulation on *csgBA* reporter activity is possibly mediated by RpoS. Indeed, although ectopic CsgD expression restored activity of the *csgBA* genes in the absence of *rpoS*, the maximal level of expression was also lower and similar to that in the complemented *hfq* strain. Furthermore, it demonstrated that expression of curli structural genes can occur independently of RpoS, which is consistent with existence of RpoD-dependent promoter upstream of the *csgBAC* operon [12, 20, 32, 64–66]. Another similarity between the *rpoS* and *hfq* strains was much reduced bimodality of curli gene expression, indicating that it is at least partly RpoS-dependent.

The impact on bimodality was even more pronounced for *ompR, mlrA* and *ihfA* genes, whose loss led to the complete abolishment of the *csgBA* activity and could not be fully compensated by ectopically induced CsgD. The corresponding regulators are known to regulate transcription of *csgD* and have multiple binding sites within the *csg* intergenic region [16, 23, 33, 65, 93]. Our results indicate that they may also directly activate transcription of curli structural genes, and that this direct regulation plays an important role in the establishment of *csgBA* bimodality, possibly by stabilizing the positive expression state. On the other hand, the negative expression state may be stabilized due to the activity of H-NS, which also has multiple binding sites within the *csg* region [16, 65]. Indeed, the *csg* intergenic region has strong inherent curvature and high AT content [94], which aids binding H-NS and IHF. This could induce sharp DNA bending [16] and, in turn, affect promoter recognition by RpoS [95] and/or accessibility of binding sites for transcription factors [96]. Indeed, it was shown that OmpR can bind simultaneously with IHF to positively regulate curli expression, whereas H-NS and IHF act in competition [16]. However, further experiments are required to elucidate how DNA curvature and the interplay between H-NS and IHF may determine the bimodality of curli expression.

A number of transcriptional factors did not show any involvement into regulation of curli fibers expression. These included LeuO, RelA, PdeL, PhoP, ZraP and QseB, but also factors that were previously described as putative curli regulators (BolA) [60] or proposed to bind to *csg* intergenic region and affect curli expression in either positive (BasR, RcdA) [58, 59], negative (BtsR, MqsR) [61, 63] or dual (RstA) [16] manner. Although this discrepancy might be due to higher complexity of curli regulation in structured communities or differences in growth conditions, our experiments make it clear that these factors are not essential for curli regulation.

In addition to transcriptional regulators, several DGCs also showed effects on curli expression in unstructured planktonic *E. coli* culture. Deletion of *dgcO* and *dgcC* resulted in decreased levels of the *csgBA* expression, dependent on the presence of DgcM/PdeR regulatory module. A previous study indicated that this effect might be mediated by the interaction of DgcO with PdeR [29], but the underlying regulatory mechanism remains to be understood. Deletion of another DGC gene, *dgcI*, had a more complex effect, reducing the number of curli-positive cells at the late stage of growth – as might be expected from a DGC gene deletion – but increasing the expression level of curli during entry into the stationary phase. This indicates that DgcI might act not only as a cyclase but also through a different regulatory mechanism.

Finally, we found that CsrD protein with degenerate diagunilate activity [40] has a positive effect on curli expression that is particularly pronounced in the absence of the DgcM/PdeR module. CsrD was reported to enable the RNaseE-mediated degradation of sRNAs CsrB and CsrC [97, 98], which are known to suppress *csgD* [89]. Moreover, interaction between CsrD and DgcM was previously observed *in vitro* [29], raising a possibility that it might be sequestered by the DgcM/PdeR complex.

Overall, this work provides a systematic analysis of regulation of curli fiber expression in unstructured planktonic *E. coli* culture and demonstrates the role of several transcription factors and chromosome-shaping proteins in bimodal activation of curli expression.

## Acknowledgements

We thank Anna Pashchenko for help with generating DGC deletion strains, Silvia González Sierra for advice on flow cytometry and Julian Pietsch for suggestions regarding data analysis and fruitful discussions. This work was supported by the Max Planck Society.

## Data availability

All data needed to evaluate the conclusions are present in the manuscript and/or supporting information.

## Conflict of interest

The authors do not declare any conflict of interest.

## Notes

### Competing Interest Statement

The authors have declared no competing interest.

### Summary of Updates

This version of the manuscript has been revised to add Supplemental Material and to correct few minor language errors.

