## Supplemental Material for "Effects of multiple cell regulators on curli gene expression in *Escherichia coli*"

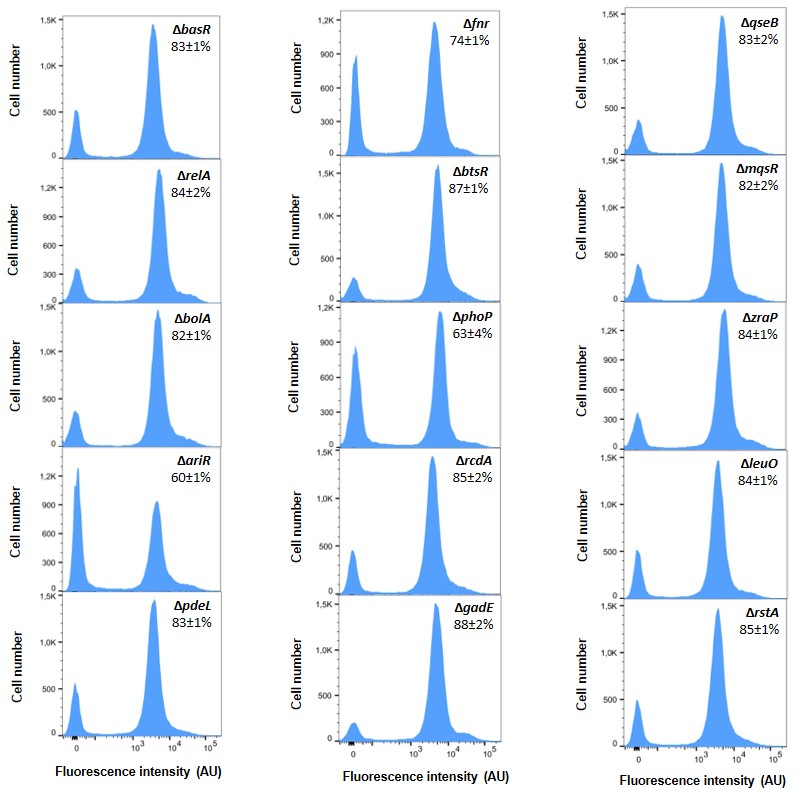
**Supplemental Figures**

**Figure S1. Curli gene expression is not affected by deletion of individual genes**. *E*. *coli* cells carrying genomic transcriptional reporter of the *csgBA* operon were grown in flasks with TB at 30˚C under constant shaking until they reach maximal *csgBA* reporter activity and then subjected to the flow cytometry analysis. Fraction of positive cells in the population (mean of at least 3 biological replicates ± SD) is indicated for each strain. Note that the scale in the *y* axes is different for individual strains to improve readability.

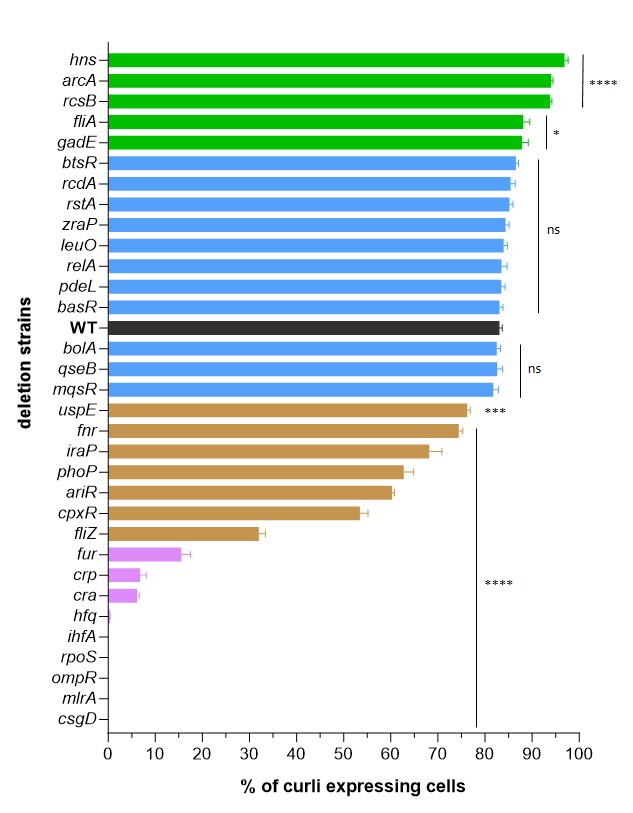

**Figure S2. Fraction of curli-expressing cells in single deletion mutants and WT.** *E*. *coli* cells carrying genomic transcriptional reporter of the *csgBA* operon were grown in flasks with TB at 30˚C under constant shaking until maximal number of curli-expressing cells is detected and then subjected to the flow cytometry analysis. WT is shown in black color and deletion strains with higher number of curli positive cells are indicated in green, with same – in blue, moderately reduced – in brown, strongly reduced – in purple and lacking – in red. Error bars indicate SEM of at least 3 biological replicates. * at p = 0.01-0.05, ** at p = 0.01-0.001, *** at p = 0.001-0.0001, **** at p < 0.0001.

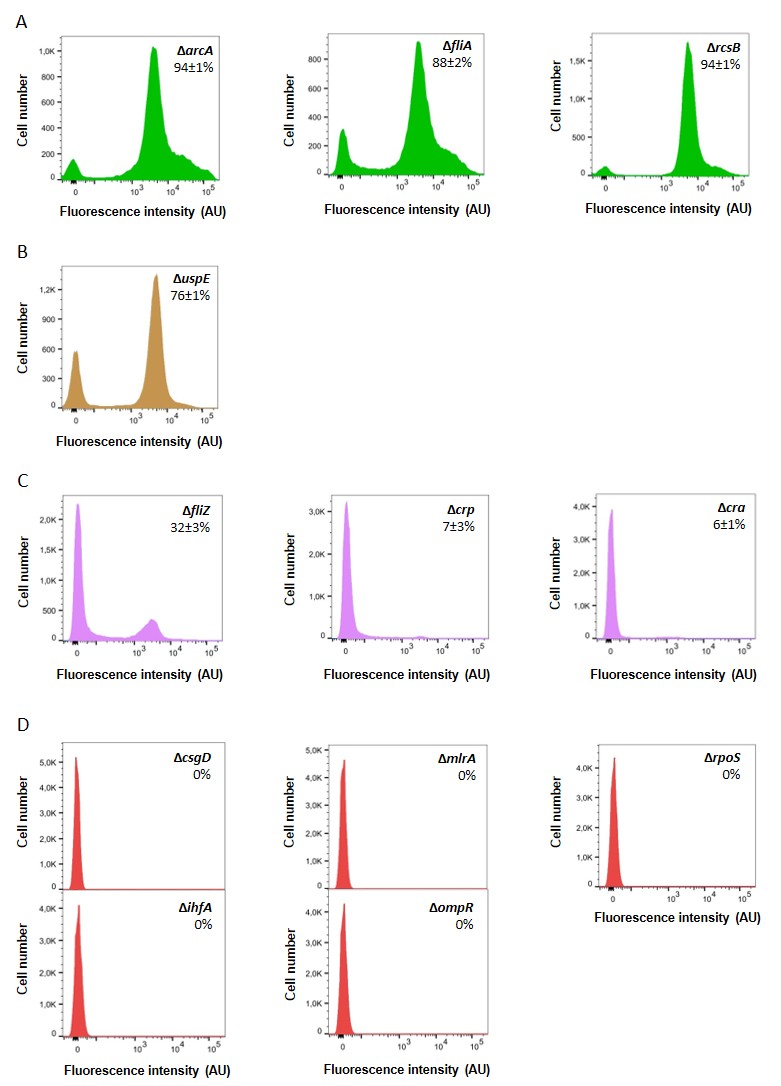

**Figure S3. Curli gene expression is disrupted upon deletion of individual genes**. *E*. *coli* cells carrying genomic transcriptional reporter of the *csgBA* operon were grown in flasks with TB at 30˚C under constant shaking until they reach maximal *csgBA* reporter levels and then subjected to the flow cytometry analysis. **(A-C)** Distribution of single-cell fluorescence levels in individual deletion strains with enhanced **(A)**, moderately reduced **(B)**, strongly impaired **(C)** and entirely abolished **(D)** curli expression. Fraction of positive cells in the population (mean of at least 3 biological replicates ± SD) is indicated for each strain. Note that the scale in the *y* axes is different for individual strains to improve readability.

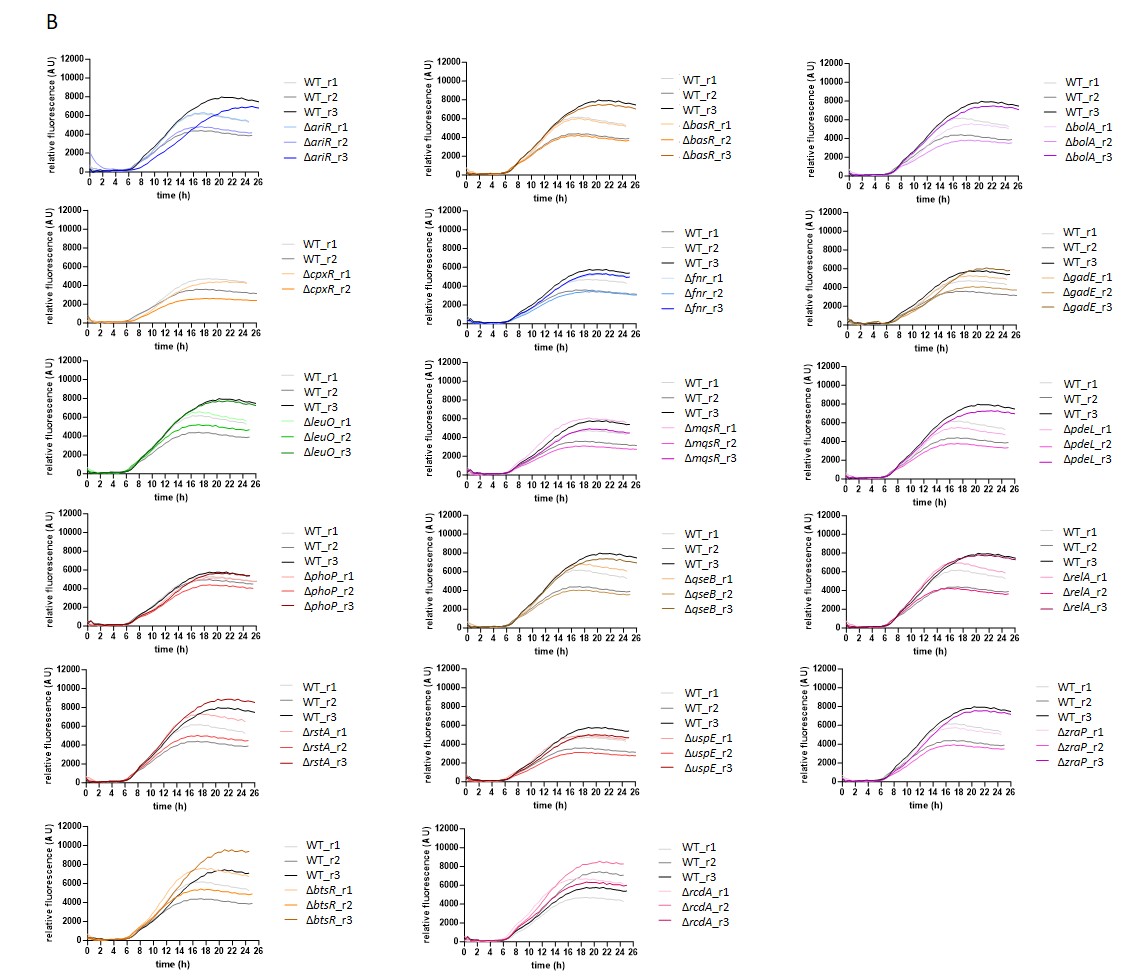

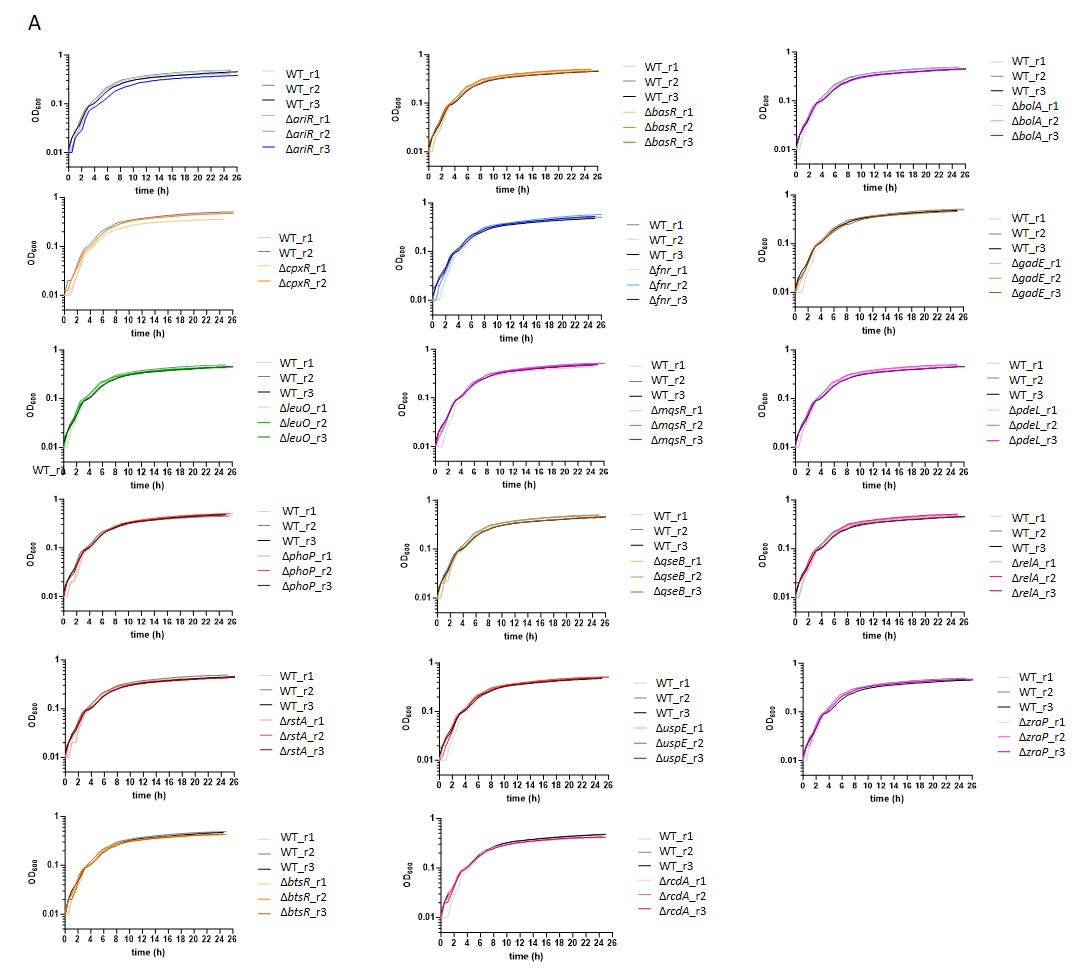

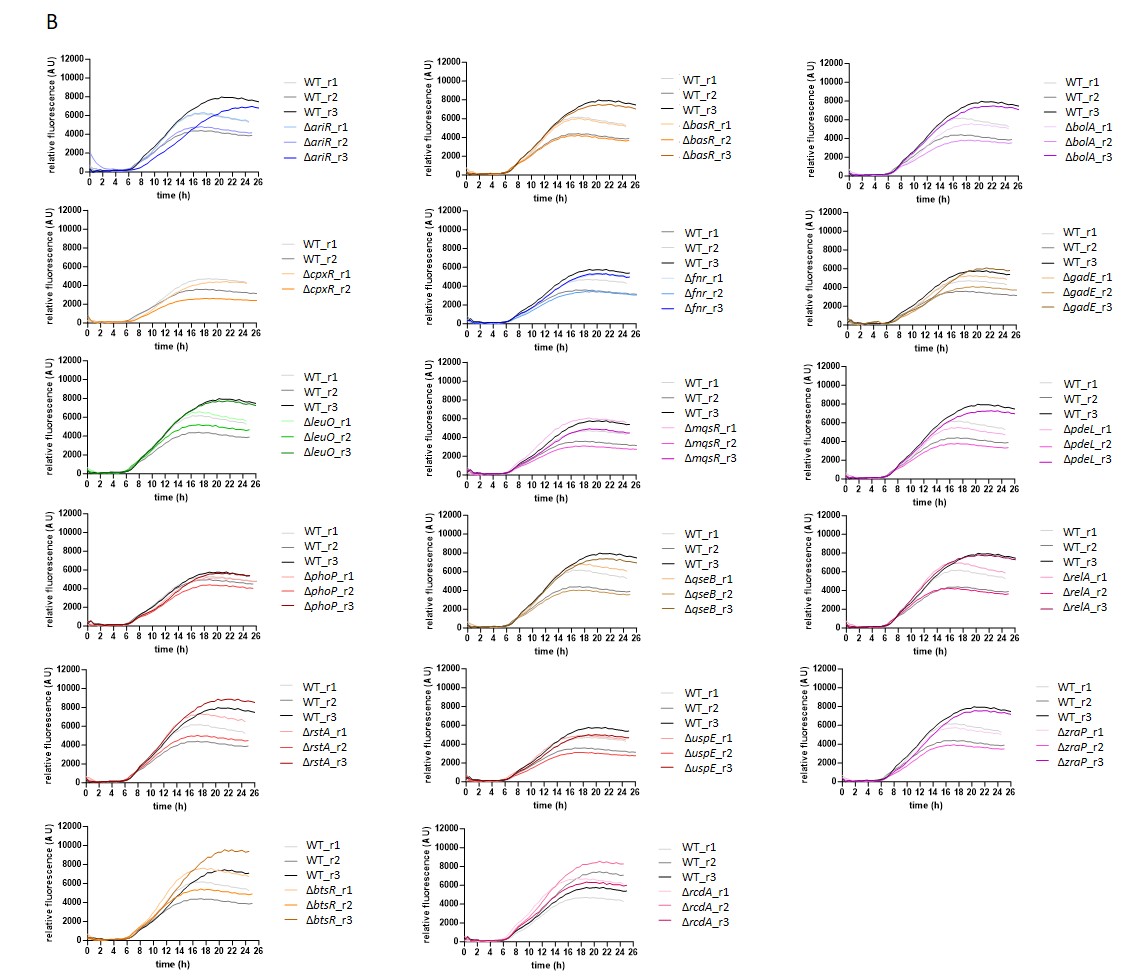

**Figure S4. Deletion of individual genes does not alter curli activation**. *E. coli* planktonic cultures carrying genomic transcriptional reporter of the *csgBAC* operon and individual gene deletions were grown in a plate reader in TB medium at 30˚C with shaking. **(A)** Optical density (OD_600_) and **(B)** relative fluorescence (absolute fluorescence/OD_600_) of WT and indicated deletion mutants during the growth in a plate reader.

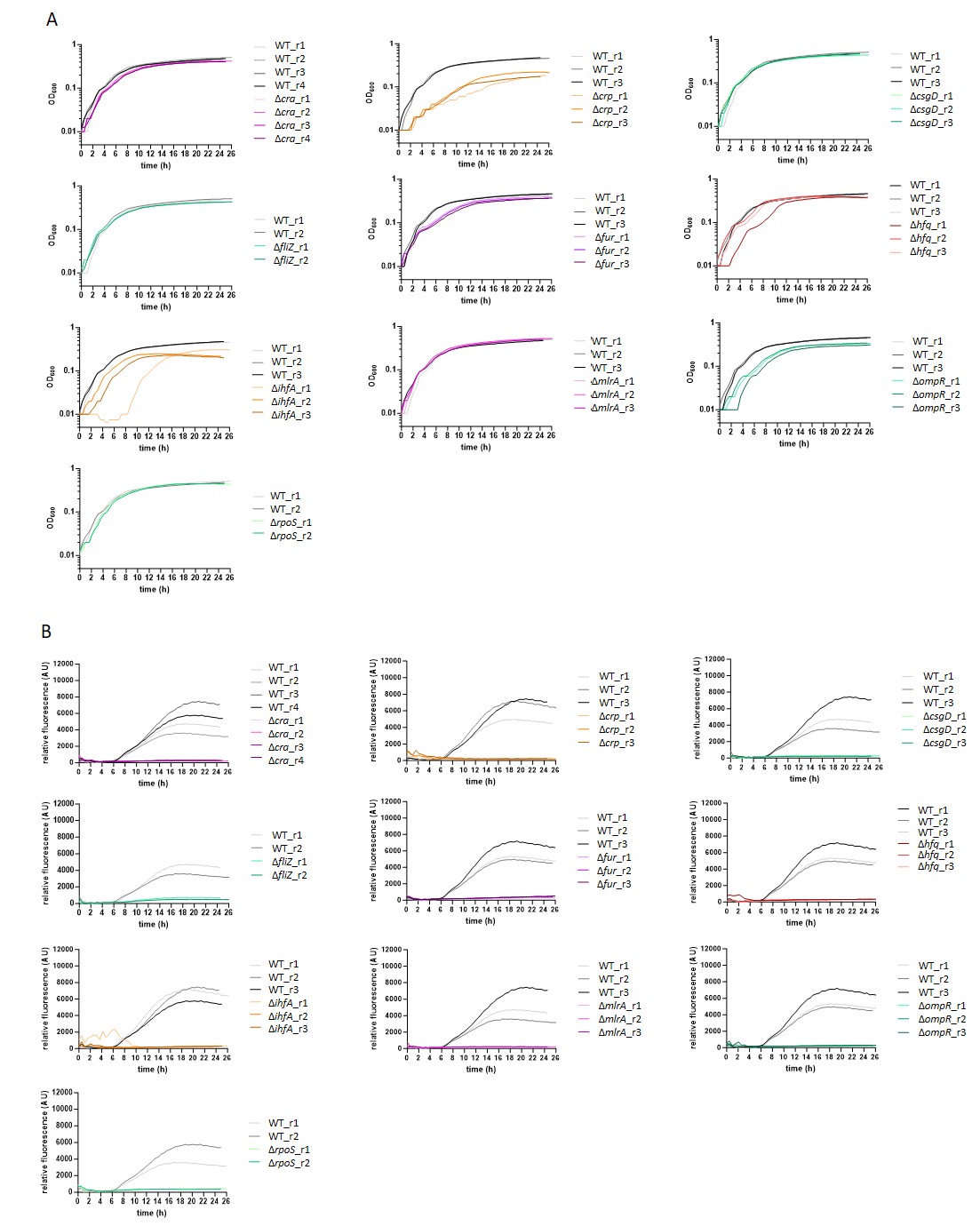

**Figure S5. Deletion of individual genes inhibits curli gene expression**. *E. coli* planktonic cultures carrying genomic transcriptional reporter of the *csgBAC* operon and individual gene deletions were grown in a plate reader in TB medium at 30˚C with shaking. **(A)** Optical density (OD_600_) and **(B)** relative fluorescence (fluorescence/OD_600_) of the WT and indicated deletion mutants during the growth in a plate reader.

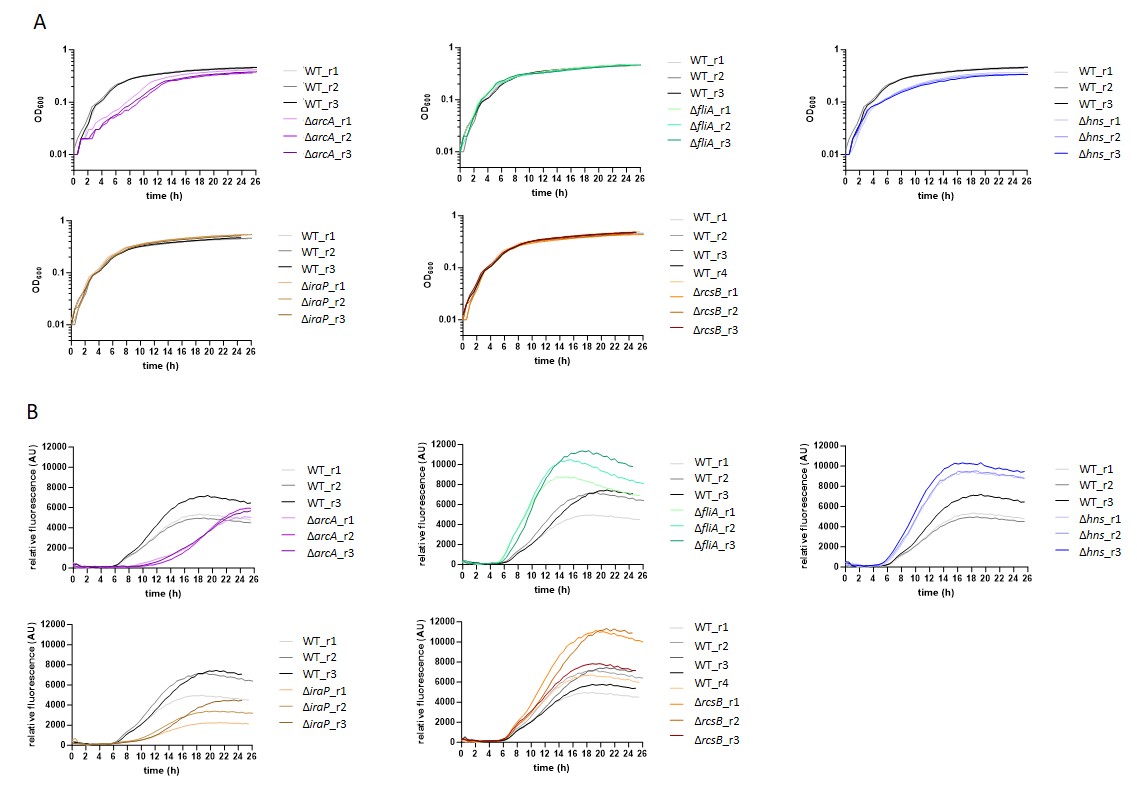

**Figure S6. Deletion of individual genes alters curli gene expression**. *E. coli* planktonic cultures carrying genomic transcriptional reporter of the *csgBAC* operon and individual gene deletions were grown in a plate reader in TB medium at 30˚C with shaking. **(A)** Optical density (OD_600_) and **(B)** relative fluorescence (absolute fluorescence/OD_600_) of WT and indicated deletion mutants during the growth in a plate reader.

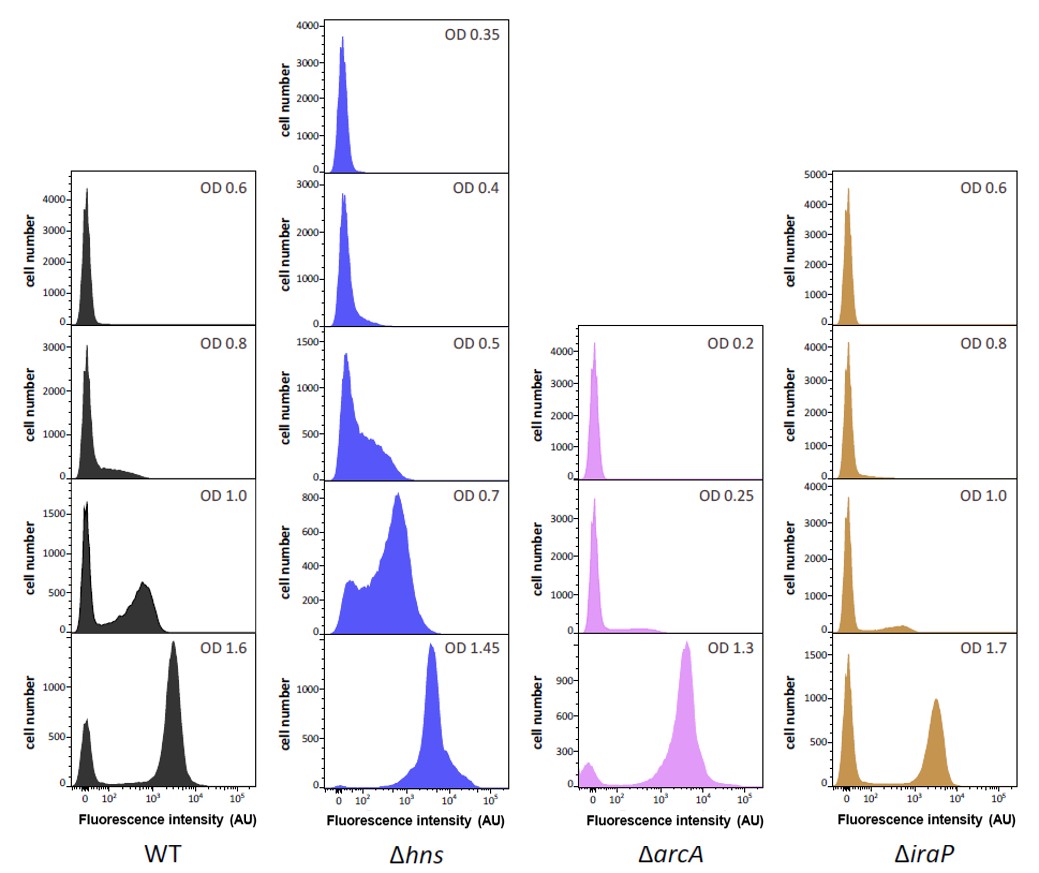

**Figure S7. Activity of the *csgBA* operon in the WT and particular deletion strains over time.**

*E*. *coli* cells carrying genomic transcriptional reporter of the *csgBA* operon were grown in flasks with TB at 30˚C under constant shaking to indicated OD_600_ and then subjected to the flow cytometry analysis. Distribution of single-cell fluorescence levels in individual deletion strains is shown. Note that the scale in the *y* axes is different for individual strains to improve readability.

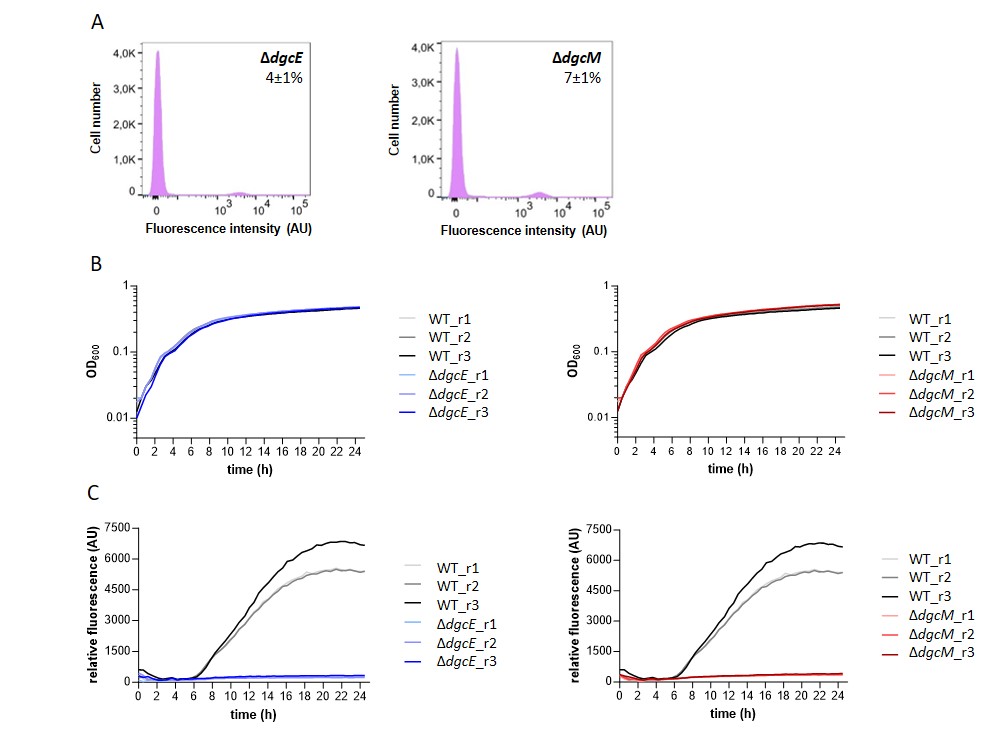

**Figure S8. Removal of individual DGCs disrupts curli gene expression.** **(A)** Distribution of single-cell fluorescence levels in deletion strains lacking individual DGCs in WT background. *E*. *coli* cells carrying genomic transcriptional reporter of the *csgBA* operon were grown in flasks with TB at 30˚C under constant shaking until they reach maximal *csgBA* reporter levels and then subjected to the flow cytometry analysis. Fraction of positive cells in the population (mean of at least 3 biological replicates ± SD) is indicated for each strain. Note that the scale in the *y* axes is different for individual strains to improve readability. **(B)** Optical density (OD_600_) and **(C)** relative fluorescence (absolute fluorescence/OD_600_) of WT and mutant strains lacking indicated DGCs during the growth in a plate reader. *E*. *coli* cells carrying genomic transcriptional reporter of the *csgBA* operon were grown in TB at 30˚C under constant shaking.

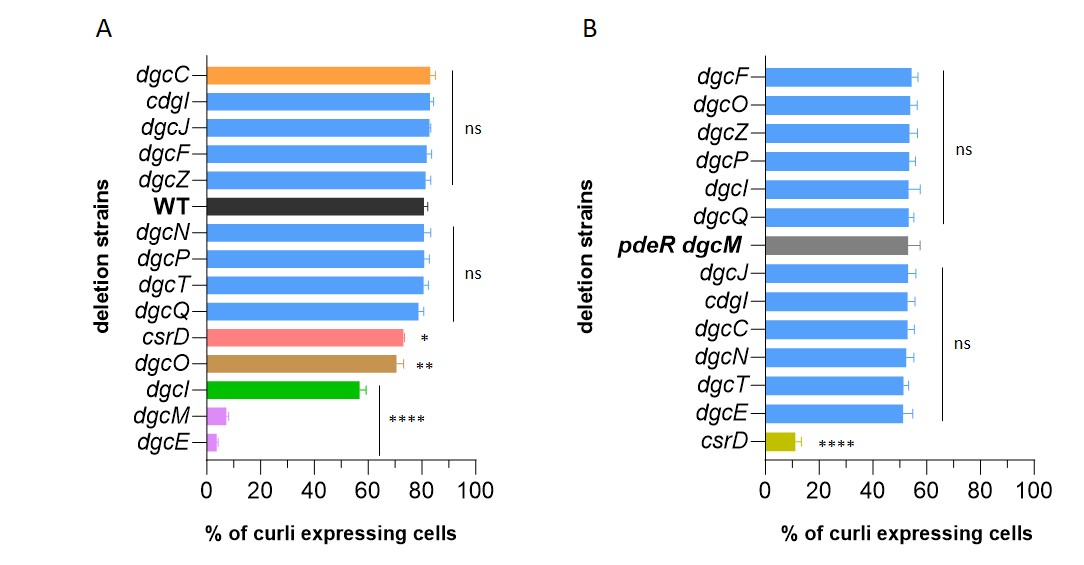

**Figure S9. Fraction of curli-expressing cells in mutants with interrupted c-di-GMP network** in WT background **(A)** and in the strain lacking local DgcM/PdeR regulatory module **(B)**. *E*. *coli* cells carrying genomic transcriptional reporter of the *csgBA* operon were grown in flasks with TB at 30˚C under constant shaking until maximal number of curli-expressing cells is detected and then subjected to the flow cytometry analysis. WT is shown in black color and deletion strains with not affected curli expression – in blue, with affected – in different colors (color code is same as in Figure 4). Error bars indicate SEM of at least 3 biological replicates. * at p = 0.01-0.05, ** at p = 0.01-0.001, *** at p = 0.001-0.0001, **** at p < 0.0001

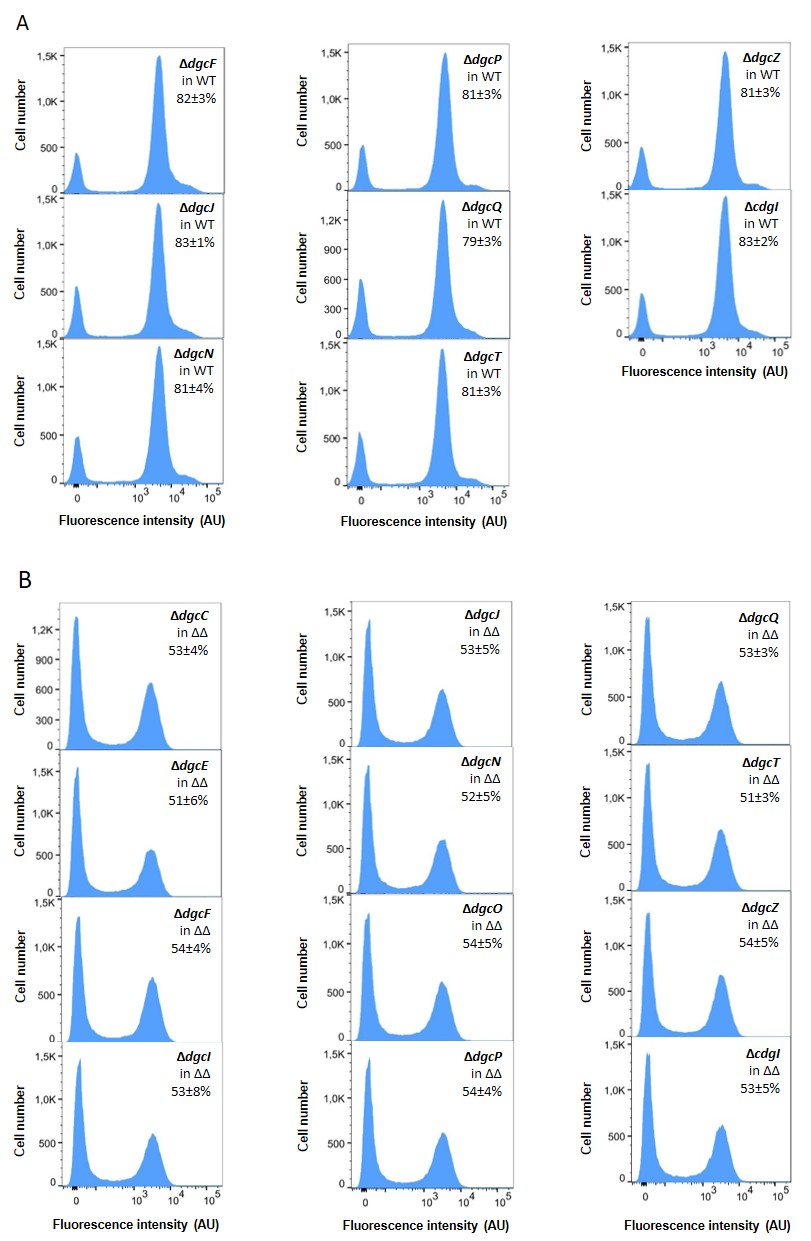

**Figure S10. Curli gene expression is not affected by removal of individual DGCs**. *E*. *coli* cells carrying genomic transcriptional reporter of the *csgBA* operon and lacking one of the indicated DGCs were grown in flasks with TB at 30˚C under constant shaking until they reach maximal *csgBA* reporter levels and then subjected to the flow cytometry analysis. Fraction of positive cells in the population (mean of at least 3 biological replicates ± SD) is indicated for each strain. Note that the scale in the *y* axes is different for individual strains to improve readability.

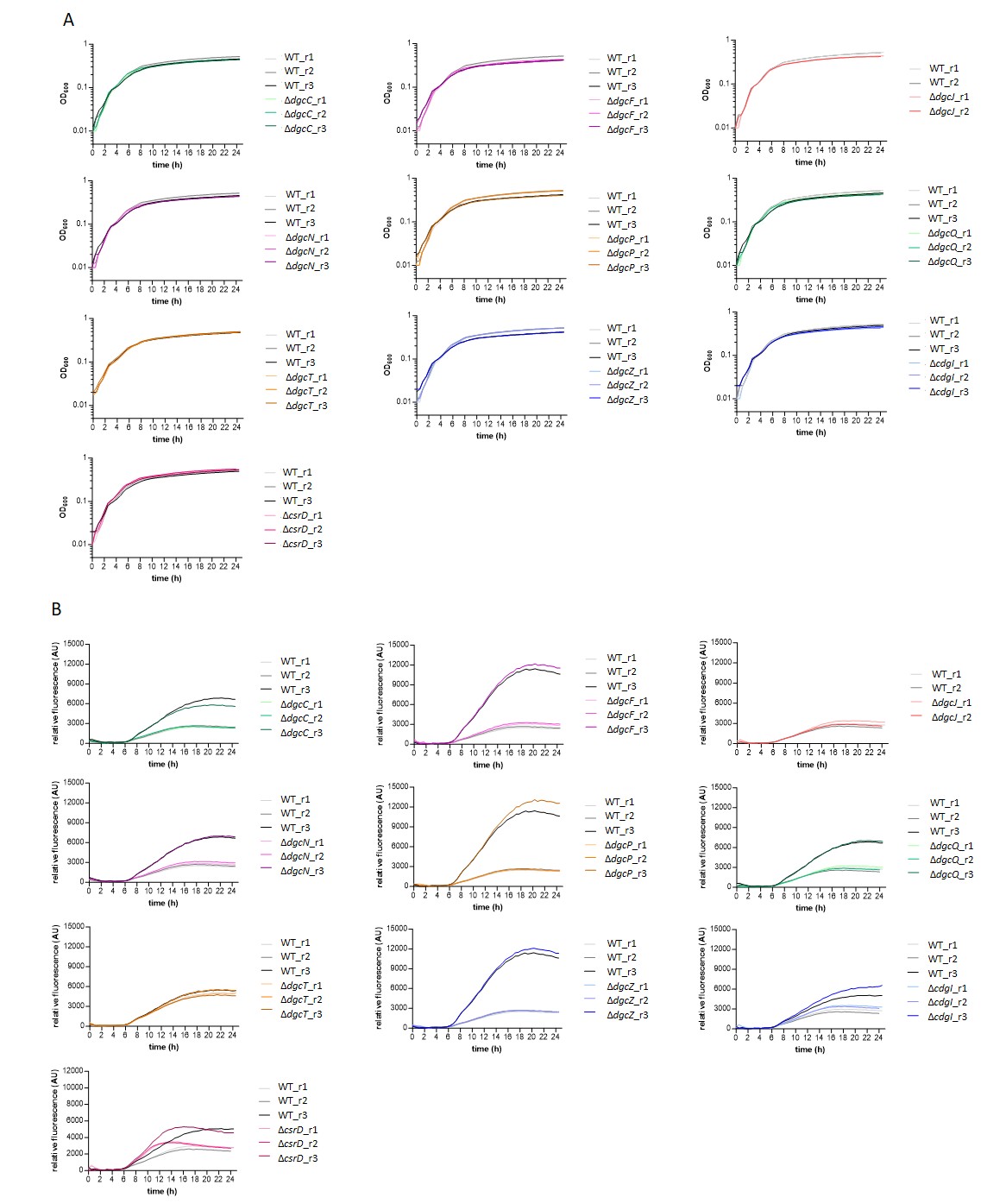

**Figure S11. Removal of individual DGCs does not alter curli activation pattern**. *E. coli* planktonic cultures carrying genomic transcriptional reporter of the *csgBAC* operon and lacking individual DGCs were grown in a plate reader in TB medium at 30˚C with shaking. **(A)** Optical density (OD_600_) and **(B)** relative fluorescence (absolute fluorescence/OD_600_) of WT and indicated deletion mutants during the growth in a plate reader.

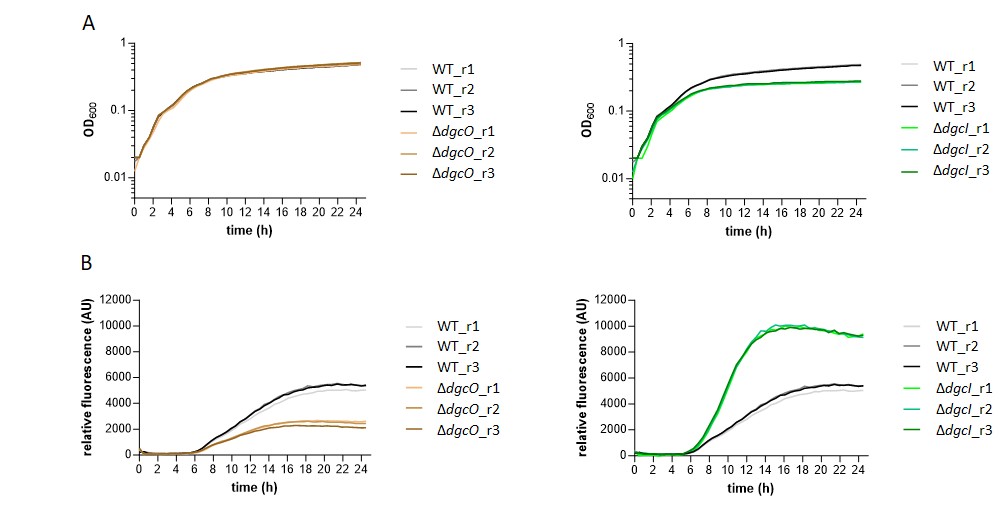

**Figure S12. Removal of certain DGCs disrupts curli gene expression.** *E. coli* planktonic cultures carrying genomic transcriptional reporter of the *csgBAC* operon and lacking individual DGCs were grown in a plate reader in TB medium at 30˚C with shaking. **(A)** Optical density (OD_600_) and **(B)** relative fluorescence (absolute fluorescence/OD_600_) of WT and indicated deletion mutants during the growth in a plate reader.

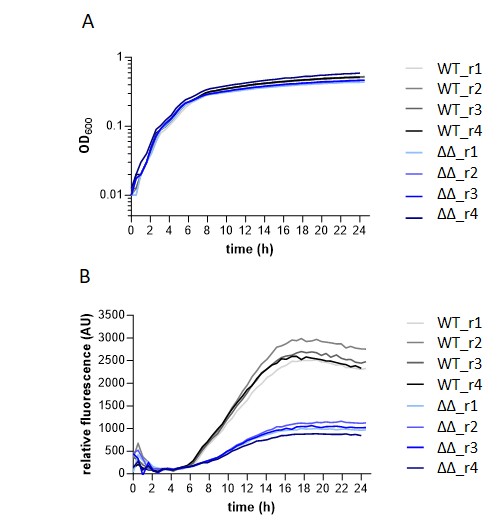

**Figure S13. Curli gene expression is disrupted upon removal of the PdeR/DgcM regulatory modulte**. *E. coli* planktonic cultures carrying genomic transcriptional reporter of the *csgBAC* operon were grown in a plate reader in TB medium at 30˚C with shaking. **(A)** Optical density (OD_600_) and **(B)** relative fluorescence (absolute fluorescence/OD_600_) of WT and strain with inactivated DgcM/PdeR regulatory module (ΔΔ) during the growth in a plate reader.

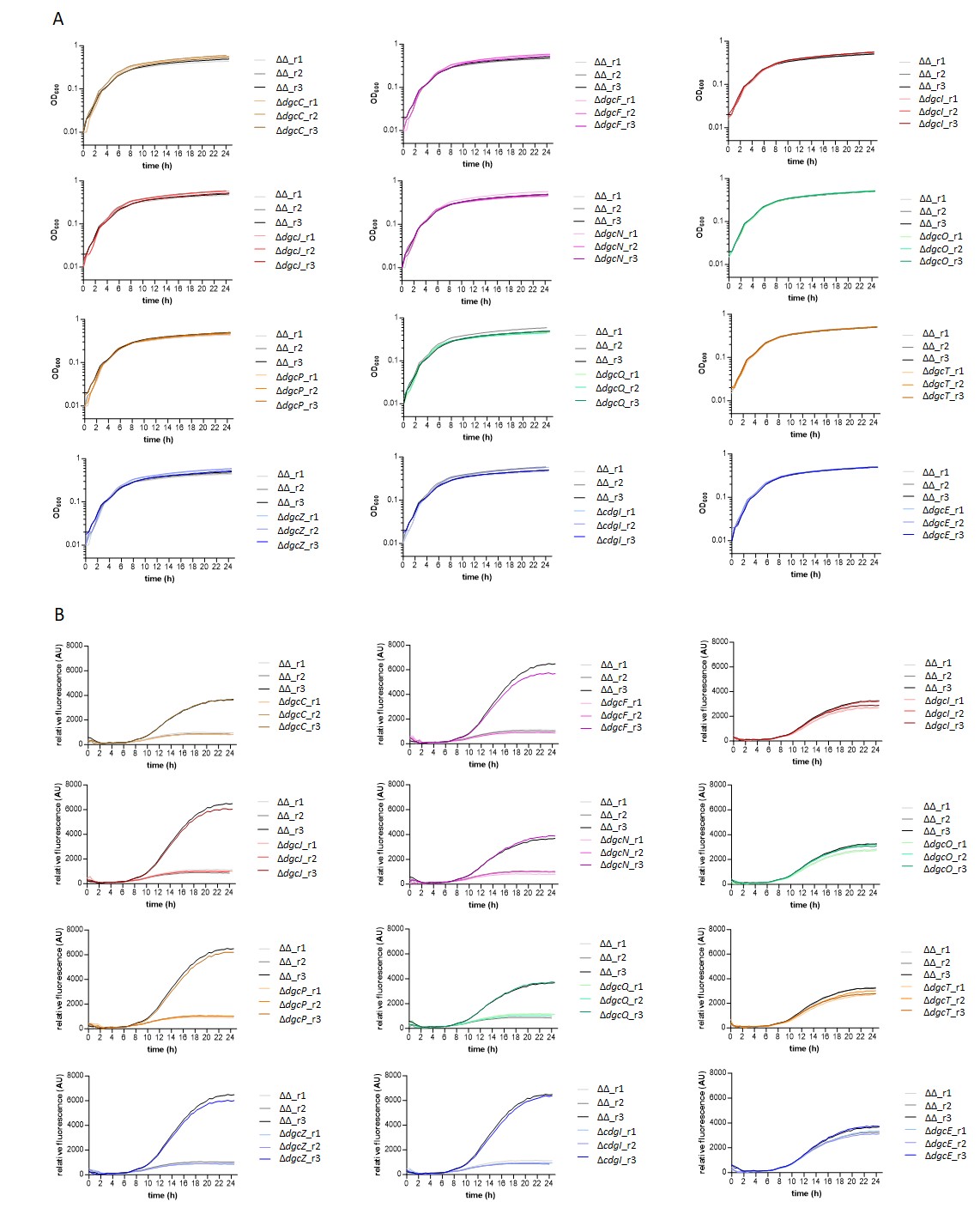

**Figure S14. Removal of individual DGCs does not affect curli gene expression in the absence of DgcM/PdeR regulatory module**. *E. coli* planktonic cultures carrying genomic transcriptional reporter of the *csgBAC* operon with inactivated DgcM/PdeR regulatory module and lacking one of the indicated DGCs were grown in a plate reader in TB medium at 30˚C with shaking. **(A)** Optical density (OD_600_) and **(B)** relative fluorescence (absolute fluorescence/OD_600_) of the strain with inactivated DgcM/PdeR regulatory module (ΔΔ) and triple deletion mutants during the growth in a plate reader.

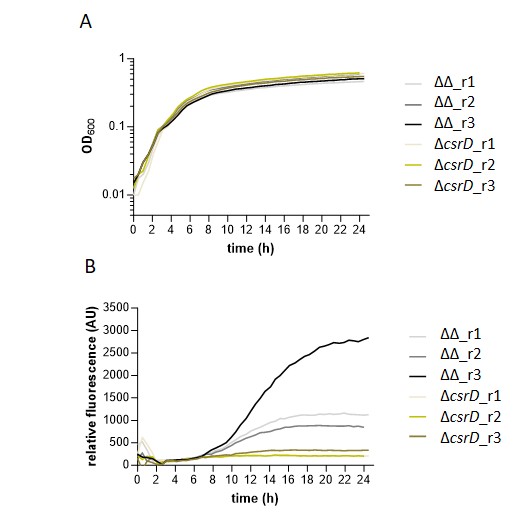

**Figure S15. Expression of curli structural genes is reduced upon deletion of *csrD* gene in the absence of local PdeR/DgcM regulatory module.** *E. coli* planktonic cultures carrying genomic transcriptional reporter of the *csgBAC* operon with inactivated DgcM/PdeR regulatory module and carrying csrD deletion were grown in a plate reader in TB medium at 30˚C with shaking. **(A)** Optical density (OD_600_) and **(B)** relative fluorescence (absolute fluorescence/OD_600_) of the strain with inactivated DgcM/PdeR regulatory module (ΔΔ) and triple deletion mutant during the growth in a plate reader.

**S1 Table. *E. coli* strains and plasmids used in this study.**

| Strains | Relevant genotype | Reference |
| --- | --- | --- |
| W3110 | W3110 derivative with functional RpoS (Km^S^) | [1] |
| VS1146 | W3110 *csgA*::*csgA-*RBS-*sfgfp* (Km^S^) | [2] |
| VS2294 | VS1146 Δ*arcA* (Km^S^) | This work |
| VS2295 | VS1146 Δ*ariR* (Km^R^) | This work |
| VS2296 | VS1146 Δ*basR* (Km^R^) | This work |
| VS2297 | VS1146 Δ*bolA* (Km^R^) | This work |
| VS2298 | VS1146 Δ*btsR* (Km^R^) | This work |
| VS2299 | VS1146 Δ*cdgI* (Km^R^) | This work |
| VS2300 | VS1146 Δ*cra* (Km^S^) | This work |
| VS2301 | VS1146 Δ*crp* (Km^S^) | This work |
| VS2302 | VS1146 Δ*cpxR* (Km^S^) | This work |
| VS2303 | VS1146 Δ*csgD* (Km^S^) | This work |
| VS2304 | VS1146 Δ*csrD* (Km^S^) | This work |
| VS2305 | VS1146 Δ*dgcC* (Km^R^) | This work |
| VS1720 | VS1146 Δ*dgcE* (Km^S^) | [3] |
| VS2306 | VS1146 Δ*dgcF* (Km^R^) | This work |
| VS2307 | VS1146 Δ*dgcI* (Km^S^) | This work |
| VS2308 | VS1146 Δ*dgcJ* (Km^R^) | This work |
| VS1257 | VS1146 Δ*dgcM* (Km^S^) | [3] |
| VS2309 | VS1146 Δ*dgcN* (Km^R^) | This work |
| VS2310 | VS1146 Δ*dgcO* (Km^S^) | This work |
| VS2311 | VS1146 Δ*dgcP* (Km^R^) | This work |
| VS2312 | VS1146 Δ*dgcQ* (Km^R^) | This work |
| VS2313 | VS1146 Δ*dgcT* (Km^R^) | This work |
| VS2314 | VS1146 Δ*dgcZ* (Km^R^) | This work |
| VS2315 | VS1146 Δ*fliA* (Km^S^) | This work |
| VS2316 | VS1146 Δ*fliZ* (Km^S^) | This work |
| VS2317 | VS1146 Δ*fnr* (Km^R^) | This work |
| VS2318 | VS1146 Δ*fur* (Km^S^) | This work |
| VS2319 | VS1146 Δ*gadE* (Km^S^) | This work |
| VS2320 | VS1146 Δ*hfq* (Km^S^) | This work |
| VS2321 | VS1146 Δ*hns* (Km^S^) | This work |
| VS1221 | VS1146 Δ*ihfA* (Km^R^) | This work |
| VS2322 | VS1146 Δ*iraP* (Km^S^) | This work |
| VS2323 | VS1146 Δ*leuO* (Km^R^) | This work |
| VS2324 | VS1146 Δ*mqsR* (Km^R^) | This work |
| VS1857 | VS1146 Δ*mlrA* (Km^S^) | [3] |
| VS2325 | VS1146 Δ*ompR* (Km^S^) | This work |
| VS2326 | VS1146 Δ*pdeL* (Km^R^) | This work |
| VS1713 | VS1146 Δ*pdeR* Δ*dgcM* (Km^S^) | [3] |
| VS2327 | VS1713 Δ*pdeR* Δ*dgcM* Δ*cdgI* (Km^R^) | This work |
| VS2328 | VS1713 Δ*pdeR* Δ*dgcM* Δ*csrD* (Km^S^) | This work |
| VS2329 | VS1713 Δ*pdeR* Δ*dgcM* Δ*dgcC* (Km^R^) | This work |
| VS1725 | VS1713 Δ*pdeR* Δ*dgcM* Δ*dgcE* (Km^R^) | This work |
| VS2330 | VS1713 Δ*pdeR* Δ*dgcM* Δ*dgcF* (Km^R^) | This work |
| VS2331 | VS1713 Δ*pdeR* Δ*dgcM* Δ*dgcI* (Km^R^) | This work |
| VS2332 | VS1713 Δ*pdeR* Δ*dgcM* Δ*dgcJ* (Km^R^) | This work |
| VS2333 | VS1713 Δ*pdeR* Δ*dgcM* Δ*dgcN* (Km^R^) | This work |
| VS2334 | VS1713 Δ*pdeR* Δ*dgcM* Δ*dgcO* (Km^R^) | This work |
| VS2335 | VS1713 Δ*pdeR* Δ*dgcM* Δ*dgcP* (Km^R^) | This work |
| VS2336 | VS1713 Δ*pdeR* Δ*dgcM* Δ*dgcQ* (Km^R^) | This work |
| VS2337 | VS1713 Δ*pdeR* Δ*dgcM* Δ*dgcT* (Km^R^) | This work |
| VS2338 | VS1713 Δ*pdeR* Δ*dgcM* Δ*dgcZ* (Km^R^) | This work |
| VS2339 | VS1146 Δ*phoP* (Km^R^) | This work |
| VS2340 | VS1146 Δ*qseB* (Km^R^) | This work |
| VS2341 | VS1146 Δ*rcdA* (Km^R^) | This work |
| VS2342 | VS1146 Δ*relA* (Km^S^) | This work |
| VS2343 | VS1146 Δ*rpoS* (Km^S^) | This work |
| VS2344 | VS1146 Δ*rscB* (Km^S^) | This work |
| VS2345 | VS1146 Δ*rstA* (Km^R^) | This work |
| VS2346 | VS1146 Δ*uspE* (Km^S^) | This work |
| VS2347 | VS1146 Δ*zraP* (Km^R^) | This work |
| Plasmids |  |  |
| pTrc99a | Expression vector; *Ptrc* promoter inducible by isopropyl-β-D-thiogalactopyranoside (IPTG); pBR ori (Ap^R)^ | [4] |
| pVS1977 | pTrc99a::*csgD* (Ap^R^) | This work |
